# Beyond Onset Timing: Longer Sound Envelope Duration Enhances Neural Representation of the Musical Beat

**DOI:** 10.64898/2026.05.12.721298

**Authors:** F Rosenzweig, C Lenoir, T Lenc, R Polak, C Huart, S Nozaradan

## Abstract

Musical rhythm is often experienced with a periodic beat, serving as a temporal reference for coordination with the rhythm. Thus far, models of beat processing have mainly relied on representing sensory inputs as patterns of onset timing, with limited consideration of other sensory features. Here, we challenge this view by showing that the internal representation of beat is affected by other temporal features of the stimulus beyond onset timing alone. We recorded electroencephalography (EEG) while participants listened to rhythmic sequences designed to elicit a beat. Across conditions, we manipulated the duration of the tones conveying the rhythms, while keeping all other parameters identical, including overall intensity, speed, and rhythmic pattern structure. Crucially, the beat periodicity was enhanced in neural activity with increased sound duration, even though the beat periodicity was not prominent in the acoustic features, thus ruling out basic sensory confounds. These results demonstrate the preferential role of longer sound durations in fostering temporal scaffolding processes that integrate fast rhythmic inputs into behavior-relevant internal structures such as the beat. More generally, our findings are compatible with a holistic processing account whereby a range of features beyond onset timing may be integrated into a neural representation of rhythm.

**Graphical Abstract: Fig. 2:** EEG was recorded while listeners heard rhythmic sequences eliciting a beat. Sound duration (sonic duty cycle) was varied across four conditions while speed, pattern, and intensity stayed constant. Beat-related EEG responses increased with longer sounds, and were enhanced in all conditions compared to auditory nerve model envelopes, which did not show prominent energy at the beat periodicity, ruling out sensory confounds. Results support holistic rhythm processing beyond onset timing alone.

## 1 INTRODUCTION

Coordinating body movement to the rhythm of music is a widespread human behavior that often entails the perception of a periodic beat. Beyond its pivotal role in music and dance, the ability to perceive and move to the beat also carries significant social ^1–3^, developmental^4,5^ and clinical implications, particularly in the assessment of higher-level brain functions for the diagnosis and rehabilitation of sensorimotor impairments ^6–12^. However, despite considerable interest in understanding the underlying processes, the mechanisms by which the brain maps periodic beats onto rhythmic inputs remain unclear.

A majority of work so far has studied this mapping by exclusively considering (i) relative timing between successive sound onsets, often described in terms of inter-onset interval (IOI) ratios ^13–19^, and (ii) the relative pattern of perceptual salience carried by these sounds (i.e., accentuation) ^20^. However, such a simplified, abstracted view of the sensory input may distort our understanding of the processes underlying beat perception. Notably, the concept of sound onset itself is far from trivial, as sound properties such as rise time have been shown to significantly alter onset perception ^21–23^ and sensorimotor synchronization to acoustic rhythms^24^. Instead, there is an emerging view that beat perception may rely on a representation that integrates features beyond patterns of onset timing alone ^25–27^.

To move beyond onset timing, prior work has also examined sound duration, quantified as the ratio between the active portion of a sound and the IOI. This ratio, hereafter referred to as the sonic duty cycle, has been shown in behavioral studies to significantly influence time perception. For instance, variations in sonic duty cycle affect the estimation of both single ^28–30^ and successive time intervals ^31,32^. Longer sonic duty cycles in isochronous sequences have been linked to a perception of event timing that is more variable across individuals and is located overall later relative to the acoustic onset ^21^, yet with more stable sensorimotor synchronization to these isochronous events ^24^.

By definition, increasing the sonic duty cycle of a fixed IOI shortens the silent interval between a tone’s offset and the subsequent onset. Studies have shown that sounds that are temporally close tend to be perceptually grouped, with silent intervals between groups being perceived as longer than their actual duration ^33^. Furthermore, two sounds that occur closer together in time than a specific just-noticeable-difference threshold are no longer perceived as different events but as merged into a single sound percept ^34,35^. Together, these findings lead to the hypothesis that, beyond influencing the encoding of onset timing, sound duration may also play a role in shaping temporal integration processes at larger timescales, which are necessary for constructing behavior-relevant internal representations such as the beat ^36^.

The influence of sonic duty cycle on rhythm and beat perception is central to real-world music production. Musicians frequently manipulate sonic duty cycles to accommodate the overall pace of the music while also adapting to external parameters such as the reverberant properties of the acoustic environment, to prevent excessive overlap or fusion between successive tones ^37^. Beyond these perceptual constraints, duty cycle manipulation is also critical for shaping the overall ’feel’ or expressivity of the music. For example, when Western popular musicians intend to give a rhythm a character such as relaxed or laid-back vs. energetic and forward-driving, they not only manipulate the timings of the produced sounds but also their duration. For instance, guitarists can convey a laid-back feeling by increasing the duration of notes ^38^. Another example comes from European art music, in which the perceived connectivity between individual sounds is itself a stylistic feature (e.g., referred to as *legato* vs. *staccato*). Overall, longer duty cycles typically produce an articulation in which one sound’s offset closely precedes the onset of the next, creating a smooth sense of flow across successive sounds. In contrast, shorter duty cycles introduce offset-onset gaps between notes, yielding a more percussive, choppy articulation. Thus, by adjusting the sonic duty cycles in conjunction with tempo and other acoustic parameters, musicians can modulate the perception of sound grouping ^39^. However, whether sonic duty cycle influences higher-level neural representations of the beat or whether, in contrast, these representations are invariant to sound duration remains debated ^40^.

Recently, advances in neuroimaging have allowed the study of neural activity related to beat processing ^25,36,41–53^. In particular, the development of new approaches combining frequency analysis with electrophysiology has offered direct insights into how the brain transforms rhythmic inputs into a neural representation that would relate to the perceived beat. Here, we took advantage of these approaches to examine whether sonic duty cycle shapes neural encoding of rhythmic inputs, and in particular, neural representation of the beat periodicity.

To this end, we recorded electroencephalography (EEG) from healthy adult participants while they listened to rhythmic sequences. To confirm the perceived beat periodicity, participants also completed a separate session in which they were asked to tap along with the same rhythmic sequences at the rate of their perceived beat. We presented four different versions of an identical rhythmic pattern looped to form long rhythmic sequences, each differing only in the sonic duty cycle of the tones conveying the rhythm while keeping all other acoustic parameters identical, including overall intensity, speed, and rhythmic pattern structure. We hypothesized that if beat perception relied solely on the pattern of sound onsets, the observed neural representation of the periodicity corresponding to the beat should remain invariant across conditions. Conversely, if the brain integrated sonic duty cycle with onset time, we should expect variations in this neural representation of the beat. This would corroborate a holistic processing account of the beat whereby different features of the rhythmic sensory inputs would contribute to the neural representation of the beat, beyond onset timing alone.

## 2 MATERIALS AND METHODS

### 2.1 Participants

Twenty-four healthy volunteers (mean age = 25.3 years, SD = 4.2 years; 17 females, 2 left-handed) participated in this study after providing written informed consent. Participants reported varying levels of musical and dance experience as amateurs (median = 2 years, range = 0–19 years). All participants had normal hearing and no history of neurological diseases. The study was approved by the Ethics Committee of the Cliniques Universitaires St-Luc, Brussels, Belgium (protocol number B403201938913).

### 2.2 Hearing Assessment

Pure-tone audiometric thresholds were assessed for each participant using TDH-39 headphones in a sound-proof booth in the Otolaryngology Department of the Cliniques Universitaires St-Luc, by trained audiologists. All participants exhibited a mean low-frequency threshold (< 500 Hz, i.e., covering the frequency range of the stimuli used in the present study) of ≤ 25 dB HL over both ears, with no asymmetry greater than 15 dB HL, and normal middle-ear pressure for all ears at the tympanometry.

### 2.3 Acoustic Stimuli

The stimuli consisted of an identical 2.4-s rhythmic pattern looped seamlessly 17 times, yielding 40.8-s duration for each sequence. The pattern comprised eight identical sound events aligned onto a grid of 12 timepoints equally distributed over the pattern duration, yielding 200-ms successive intervals (Fig. 1). The sound events consisted in pure tones with 10-ms linear onset/offset ramps and a carrier frequency of 300 Hz. This low carrier frequency was chosen to optimize beat-related neural responses, based on prior evidence that rhythms conveyed with carrier frequencies in similar low-frequency range elicit larger beat-related neural responses ^25^. Critically for the goal of the experiment, the duration of the constituent sound events was systematically manipulated across four conditions (50 ms, 100 ms, 150 ms and 200 ms), while keeping other parameters constant, including the timepoint grid and rhythmic pattern structure. Thus, the stimulus sequences across conditions differed in the ratio of active sound time to the length of the individual grid interval. In other words, the ratio of duty cycle to grid interval (200 ms) increased linearly across conditions: 25%, 50%, 75% and 100%, respectively (Fig. 1).

**Figure 1.**
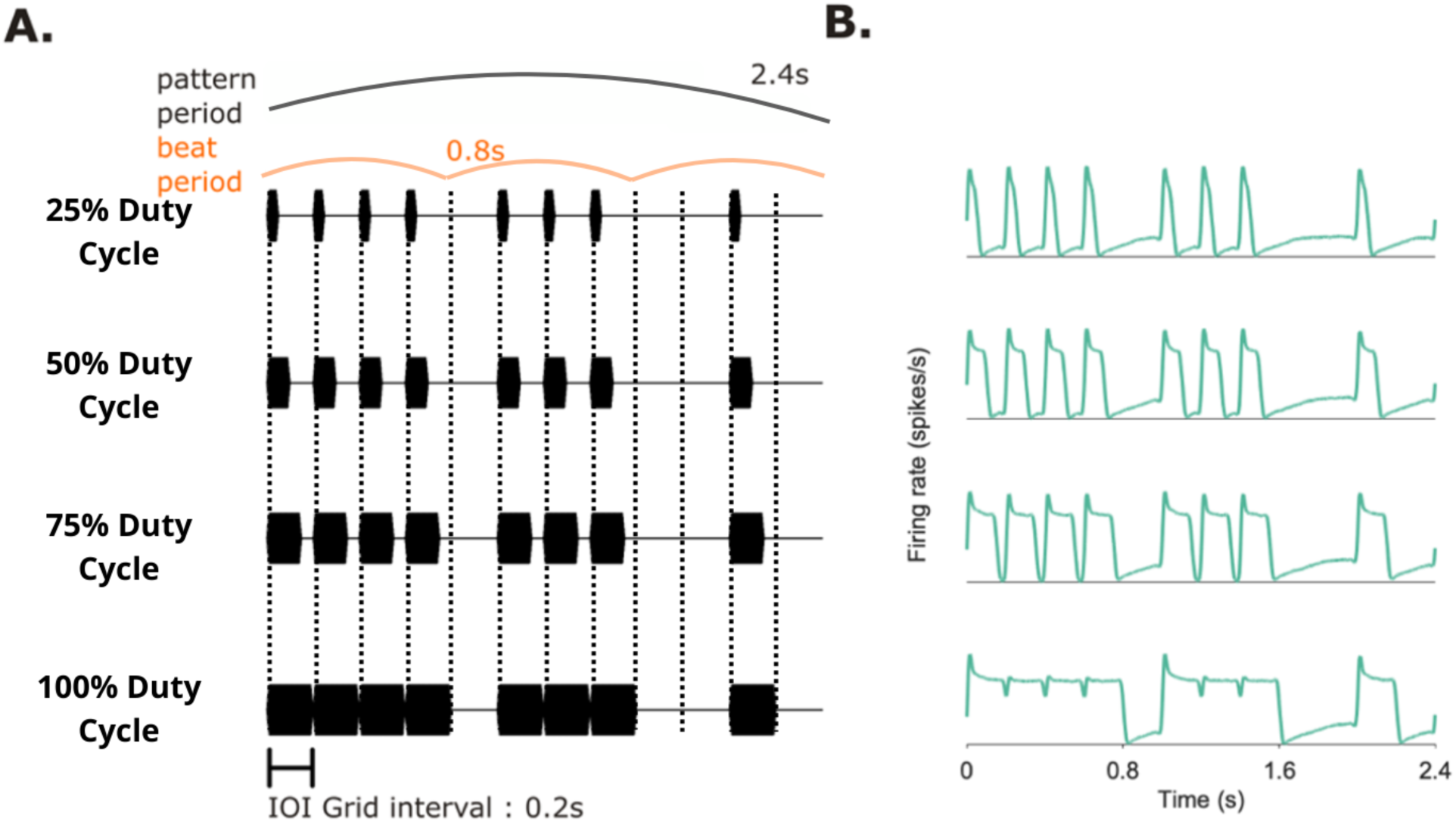
Stimuli using the same rhythm, with four conditions manipulating the sound duration (duty cycle). A) Time-domain representation of the four acoustic stimuli, each row showing one cycle of the 2.4 s rhythmic pattern repeated to form the 40.8 s sequences. The sequences were composed of tones with either 50 ms, 100 ms, 150 ms, or 200 ms duration (from top to bottom), spaced on a 200 ms inter-onset-interval (IOI) grid (dotted vertical lines), thus corresponding to 25%, 50%, 75% and 100% sonic duty cycle, respectively. The onset times of sound events making up the rhythmic pattern were fixed across all sequences, and the perceived beat period was found to correspond to 800 ms across conditions (illustrated in orange). B) Time-Domain representation of the auditory nerve model output (one 2.4 s cycle depicted for each sequence; mean firing rate across all simulated auditory nerve fibers in spikes per second).

The rhythmic pattern used here is known to elicit beat perception in listeners enculturated in Western music, with a convergent beat periodicity corresponding to four grid intervals (i.e., 4 x 200 ms = 800 ms) ^25,36,50,51^. This rhythmic pattern has also been shown to elicit prominent neural responses at frequencies corresponding to this periodicity and its harmonics ^25,36,50,54–56^. Crucially, the periodicity corresponding to this perceived beat is not salient in the acoustic features of the rhythmic pattern ^19^, which can be quantified by analyzing its envelope modulations (see also section auditory nerve model analysis, below). In this respect, this rhythm can thus be qualified as “syncopated” or “weakly-periodic” ^25,36^. These stimulus characteristics are important in the current study design as they allow us to control for lower-level sensory confounds by ruling out that prominent neural responses elicited at the beat periodicity may result from neural tracking of prominent features of the envelope ^57–59^.

To match the overall intensity of the stimuli across conditions, magnitudes were normalized using the root mean square (RMS) level of each entire sequence. This was achieved by scaling each sequence’s amplitude relative to the 100% duty cycle sequence which contained the largest total energy (RMS = 0.56), using the following factors: 2.26 (25%), 1.47 (50%) and 1.17 (75%). Finally, sound intensity was set to approximately 70 dB SPL to ensure comfortable and consistent presentation levels across participants.

The acoustic stimuli were created using MATLAB R2020a (The MathWorks, Natick, USA) and presented binaurally through insert earphones (ER 2, Etymotic Research, Elk Grove Village, IL, USA; with air-conducted sound delivered from the clavicle level to minimize magnetic interferences), through a Fireface UC audio interface (RME Audio, Haimhausen, Germany; sampling frequency = 44100 Hz). Stimuli were delivered using the Psychtoolbox extension in MATLAB (version 3.0.16, Brainard 1997).

### 2.4 Experimental design

Each of the four sequences was presented in a separate block consisting of ten EEG "static" trials, followed by four "tapping" trials of the same 40.8-s sequence. The order of the blocks was counterbalanced across participants to minimize bias in the collected data due to non-specific effects (such as learning and fatigue) over the course of the experiment. Prior to the experiment, participants completed a training session to ensure they understood the instructions of the tapping task (see Supplementary Materials for details).

During the EEG "static" trials, a sham task ensured attention was allocated to the temporal aspects of the sequences (instruction: "Report at the end of each trial if you perceived very slight variations in the tempo of the sequence”). After each trial, participants reported whether they had detected any tempo variation.

In the “tapping” trials, participants were instructed to tap with their preferred index finger on a custom-built response box, along with the periodic beat they perceived, as soon as the sequences started and throughout the whole sequence duration (see Supplementary Materials for details). These tapping trials aimed to establish a group-level estimate of the perceived beat period ^60^, and empirically confirm expectations based on previous research ^25,36,50,51^. Such investigation of the perceived beat periodicity is critically informative for the interpretation of the respective EEG signals across conditions, as detailed below.

### 2.5 EEG Acquisition and Preprocessing

Participants were comfortably seated with their head resting on a support. They were instructed to relax, minimize body movements, and fixate a cross placed approximately 1.5 meters in front of them. EEG data were recorded using a Biosemi Active-Two system with 64 sintered Ag-AgCl pin-type active electrodes positioned on the scalp according to the International 10/20 system (Biosemi, The Netherlands). The signal was referenced to the CMS (Common Mode Sense), with electrode offsets kept below ±30 mV. Two additional flat-type active electrodes were placed on the mastoids, with offsets kept below ±50 mV. Data were digitized at a 1024 Hz sampling rate.

EEG preprocessing was performed using the open-source Letswave toolbox versions 6 and 7 (https://www.letswave.org/) under MATLAB R2020a. The continuous time-domain signals obtained for each participant, condition and trial were band-pass filtered from 0.1 to 45 Hz using a 4^th^-order Butterworth filter to remove very slow drifts and higher frequencies irrelevant to the aim of the current study. The data were then segmented from -1 to +40.8 s relative to the trial onset, thus encompassing a 1-s time window before stimulus presentation followed by the response to the whole duration of each sequence. Epochs were visually inspected to detect noisy electrodes. In four participants, one electrode was interpolated using the six closest electrodes. Due to technical issues, one participant had nine epochs instead of ten in the 75% condition.

Eyeblink artifacts were removed using an independent component analysis (ICA) method ^61^ as implemented by the infomax algorithm ^62,63^. ICA matrices were computed from an alternative preprocessing pipeline in which the original continuous time-domain datasets were high-pass filtered at 1 Hz (4^th^-order Butterworth filter) for optimal artifact classification accuracy ^64,65^. The data were then segmented from -1 to +40.8 s relative to trial onset to include pre-stimulus baseline activity, reducing the potential ICA bias towards response-related activity ^66^. Matrices were computed on a merged dataset combining all four duty cycle conditions and all epochs (i.e., data recorded from each trial) per participant, with the number of components (IC) constrained to 63 minus the number of interpolated channels for each participant ^67^. The matrices were then applied to the original individual band-pass filtered datasets to remove artifacts related to eye blinks and movements (1 IC in 19 participants, 2 ICs in 5 participants). The epochs were then re-segmented from 0 to +40.8 s to discard pre-stimulus activity, and were re-referenced to the averaged mastoid channels to improve signal-to-noise ratio of neural responses to acoustic stimuli ^68^.

### 2.6 EEG Analysis

For each participant and duty cycle condition, preprocessed time-domain data were averaged across epochs to increase signal-to-noise ratio of the responses time-locked across trials ^69^. The resulting averaged epochs were then transformed into the frequency domain using a fast Fourier transform (FFT), yielding a normalized magnitude spectrum (in μV) over the range of 0 to 512 Hz with a frequency resolution of 0.0245 Hz (1/40.8 s).

A baseline correction was applied to each obtained spectrum to minimize the contribution of local variations of noise inherent to EEG recordings. Baseline was defined as the mean magnitude at -2 to -10 and +2 to +10 frequency bins relative to each frequency bin over the spectrum ^50,70,71^. Noise-subtracted spectra were then averaged over a pool of fronto-central electrodes (F1, FC1, C1, F2, FC2, C2, Fz, FCz, Cz), where responses to the acoustic rhythms were expected to be largest based on previous research ^25,50,51,72–74^.

From the obtained noise-subtracted, channel-averaged EEG spectra, we then measured the magnitude at frequencies of interest determined as the frequencies corresponding to the repetition rate of the rhythmic pattern and its harmonics (i.e., 1/2.4 s = 0.416 Hz, x 2, x 3, x 4, etc.) extending up to the repetition rate of individual events within the pattern (i.e., 0.416 Hz x 12 = 5 Hz, corresponding to the 1/0.2 s IOI). This set of frequencies is expected to capture the major part of the overall neural responses to the repeated rhythmic patterns, as demonstrated in previous EEG work using similar rhythmic stimuli and frequency-tagging analysis framework _25,36,49,50,73,75–77._

#### 2.6.1 Measuring prominence of the beat period in brain responses: Magnitude spectrum-based analysis

From this set, the first two frequencies were excluded from further analyses as they were located within the frequency range most affected by 1/f-like background noise (< 1 Hz) ^36,68,78^. The twelfth frequency (5 Hz) corresponding to the rate of individual sonic events was also discarded, as it primarily reflects the neural response elicited by the single sounds composing the rhythmic pattern rather than beat periodicity (as demonstrated in Fig. S2 of ^49^). This yielded a total of nine frequencies of interest that were further classified as beat-related and beat-unrelated. A frequency was considered as beat-related when corresponding to the frequency of the beat and harmonics, or beat-unrelated otherwise. The frequency of the beat was inferred from the group-level main tapping beat period, which converged toward four grid intervals (4 x 0.2 = 0.8 s), consistent with prior studies using the same pattern, tempo and participant recruitment criteria (see Supplementary Materials for full details of the tapping analysis and results) ^25,51,54,77,79^.

The magnitudes at all frequencies of interest were extracted and standardized as z-scores, in order to capture their *relative* prominence irrespective of the original unit and scale ^49^. The z-score at each frequency of interest therefore expressed their relative prominence with respect to the entire set of frequencies of interest. To quantify the prominence of beat periodicity in each sequence, individual beat-related frequency z-scores were averaged, yielding a mean beat-related z-score per participant and duty cycle condition. Positive values would thus reflect prominent beat periodicity in the EEG responses.

#### 2.6.2 Measuring beat prominence in brain responses: Autocorrelation-based analysis

Although the magnitude spectrum provides a highly sensitive measure of the prominence of beat periodicity, it can also be influenced by beat-unspecific effects such as the shape of the brain responses to individual sound events within the stimulus sequence. In the present study, the individual sound envelope durations varied across conditions, making it particularly important to determine whether any differences in beat-related EEG responses reflected envelope manipulations rather than changes in internal representation of the beat period per se. To this end, a complementary approach using autocorrelation was applied. First, to minimize contamination with noise in further calculating the autocorrelation, a two-step noise-correction method was applied to each epoch, as described in [^49^]. A first step consisted in subtracting from the data the 1/f-like noise baseline. To do so, the spectrum of the 1/f-like noise baseline was estimated using irregular resampling with scaling factors between 1.1 and 2 in steps of 0.05 (as implemented in the IRASA package; see ^80^). The data were subsequently transformed into the frequency domain using FFT and the 1/f-like noise estimate was subtracted from the resulting complex spectrum. A second step to further reduce the contribution of noise consisted in setting to zero all complex Fourier coefficients at frequency bins that did not correspond to the frequencies of interest (i.e., harmonics of the rhythm repetition rate). Finally, circular autocorrelation was obtained from the noise-corrected spectra.

The resulting autocorrelation function reflects the similarity of the signal with a lagged version of itself, across a range of lags from 0 s to half of its duration, here corresponding to half the total duration of the rhythmic sequence (i.e. 40.8 s / 2 = 20.4 s). From this autocorrelation function, we extracted values at lags of interest corresponding to beat periodicities (beat-related lags: 0.8 s and multiples thereof) and beat-unrelated periodicities determined as multiple of the grid interval that did not correspond to the beat periodicity and its multiples (beat-unrelated lags: 0.6 s, 1 s, 1.4 s and 1.8 s and multiples). Lags that overlapped between the two categories were excluded. After normalizing the autocorrelation coefficients across the whole set of lags using z-scoring, the periodicity of the response at the rate of the beat was quantified by averaging the coefficients across beat-related lags, yielding a beat-related z-score that provides an index of periodicity that is invariant to the shape of the recurring signal. In other words, autocorrelation captures how robustly the response repeats itself at the rate of the beat, regardless of the specific shape within the recurrent period (for further methodological details, see [^36^ ]; see also [^49^]).

#### 2.6.3 Confirming the frequency bandwidth of brain responses across duty cycle conditions

We also evaluated whether the set of frequencies of interest, as determined above based on previous EEG studies using similar rhythmic stimuli, covered a frequency range (< 5 Hz) that was adequately matching the actual spectral distribution of the EEG responses obtained here across all duty cycle conditions. Indeed, the spectral distribution of the response might shift toward higher frequencies as duty cycle varied.

To test this, we summed for each condition and individual noise-subtracted spectrum the magnitudes of all harmonics of the pattern repetition rate (i.e., 1/2.4 s = 0.416 Hz, x 2, x 3, x 4, etc.) up to 45 Hz (i.e., the upper limit of the EEG bandpass filter). Magnitudes were also summed within successive 5-Hz-wide bands from 0 to 45 Hz, thus yielding nine frequency bands. For each participant and condition, the contribution of each band was quantified as the ratio of the band-specific summed magnitude to the total summed magnitude across all harmonics up to 45 Hz. Group-level band contributions were then compared across conditions. Note that this 5-Hz band approach was carried out for two reasons. Firstly, it ensured that each sub-band contained the same proportion of beat-related and -unrelated frequencies. Secondly, considering groups of harmonics at a time reduces the risk of missing a substantial portion of signal’s bandwidth due to a single weak harmonic near the noise floor. Indeed, because the rhythmic pattern making up the stimulus sequence was based on a grid of twelve equal intervals, its envelope spectrum was expected to exhibit substantial repetition every 12 harmonics (equivalent to 5 Hz in this case), with some harmonics having magnitudes close to zero (e.g., the sixth harmonic) ^49^.

### 2.7 Auditory Nerve Model Analysis

Finally, to assess whether the EEG responses could be explained by low-level auditory representations of the stimuli, the four stimulus sequences were analyzed using an auditory nerve model from the UR_EAR_2020b toolbox in MATLAB ^81^.

The UR_EAR_2020b toolbox integrates several computational models simulating neural responses to auditory stimuli at various levels of the auditory pathway by applying sequential signal processing steps to acoustic inputs. This provides insight into the nonlinearities introduced by early auditory processing and a baseline from which higher-level contributions could be disentangled. The auditory nerve model used in the current study was developed by ^82^ as an extension of ^83^. The model comprises a middle ear linear bandpass filter followed by cochlear nonlinear filtering by chirp filter banks dynamically modulated by an outer hair cell model. An inner hair cell model rectifies and lowpass-filters the signal, thereby extracting its envelope. The last step simulates auditory nerve synaptic properties by accounting for firing rate adaptation. The final output of the model thus consisted of N firing rate time series (in spikes per second) representing the neural activity of individual auditory nerve fibers.

The pure-tone thresholds (in dB HL) used for the model were obtained by averaging participants’ audiograms for both ears. The analysis was divided over 64 cochlear channels with center frequencies logarithmically spaced between 125 Hz and 8000 Hz. Cochlear tuning function estimates were derived from otoacoustic and behavioral measurements by Shera et al. (2002). For each channel, a simulation of 51 auditory nerve fibers was computed, with 16% low, 23% medium, and 61% of high-spontaneous-rate fibers. These proportions and thresholds were derived from data described in [^85^]. The simulated firing rates were then summed across all fibers and channels, resulting in an analytical mean rate across fibers in response to each sequence (Fig. 1). The output for each sequence was then analyzed in the frequency domain using the same analysis steps as for the EEG responses, for comparability, thus yielding a measure of the prominence of beat periodicity across the four auditory nerve model sequences. Importantly, the z-score standardization allowed direct comparison of beat-related prominence between the auditory nerve model outputs and EEG responses, thereby capturing the degree of transformation between low-level representations and higher-level brain responses.

The frequency bandwidth of the auditory nerve model outputs was also characterized using the same approach as for the EEG responses. Specifically, the magnitudes of harmonics of the rhythmic pattern were summed within the same successive 5-Hz frequency bands as in the EEG analyses, yielding proportions of the total spectral magnitude across all harmonics up to 45 Hz.

### 2.8 Statistical Analysis

Analyses were conducted in R (version 4.3.2; R Core Team, 2023). Beat-related z-scores from the magnitude spectrum- and autocorrelation-based analyses were examined using separate linear mixed-effects models (LMMs) from the ‘lme4’ package (version 1.1-35.1) for each analysis, with participant included as a random intercept. The subject-level proportion of EEG magnitude below 5 Hz (relative to 0–45 Hz) was examined using a separate LMM. For all LMMs, model assumptions (normality and homogeneity of variance) were checked using quantile-quantile and residual plots. Duty cycle was entered either as a categorical fixed effect (for comparison across conditions) or as a continuous predictor (for the bandwidth analysis).

For the magnitude spectrum-based analysis, a separate LMM tested the relationship between auditory nerve model and EEG beat-related z-scores, with auditory nerve model z-scores entered as a continuous predictor. Fixed effects were assessed using Type II Wald F-tests with Kenward–Roger degrees of freedom as implemented in the ‘car’ package (version 3.0-5). Estimated marginal means (EMMs) for each duty cycle condition were computed using the ‘emmeans’ package. Where applicable, post-hoc pairwise comparisons were performed with Bonferroni correction for multiple comparisons. Moreover, polynomial contrasts were used to test whether EEG beat-related z-scores changed monotonically with duty cycle.

Finally, each condition’s EMM was tested against the corresponding auditory nerve model beat-related z-score, using one-sample t-tests, to assess for periodization of neural responses beyond lower-level auditory representations. Partial eta-squared (η_p_²) values were computed from the Type II F-tests as a measure of effect-size.

## 3 RESULTS

### 3.1 Magnitude spectrum-based analysis

In all four conditions, we obtained EEG responses at beat-related and -unrelated frequencies with a fronto-central topographical distribution (Fig. 2), consistent with previous work ^25,55,72^.

**Figure 2.**
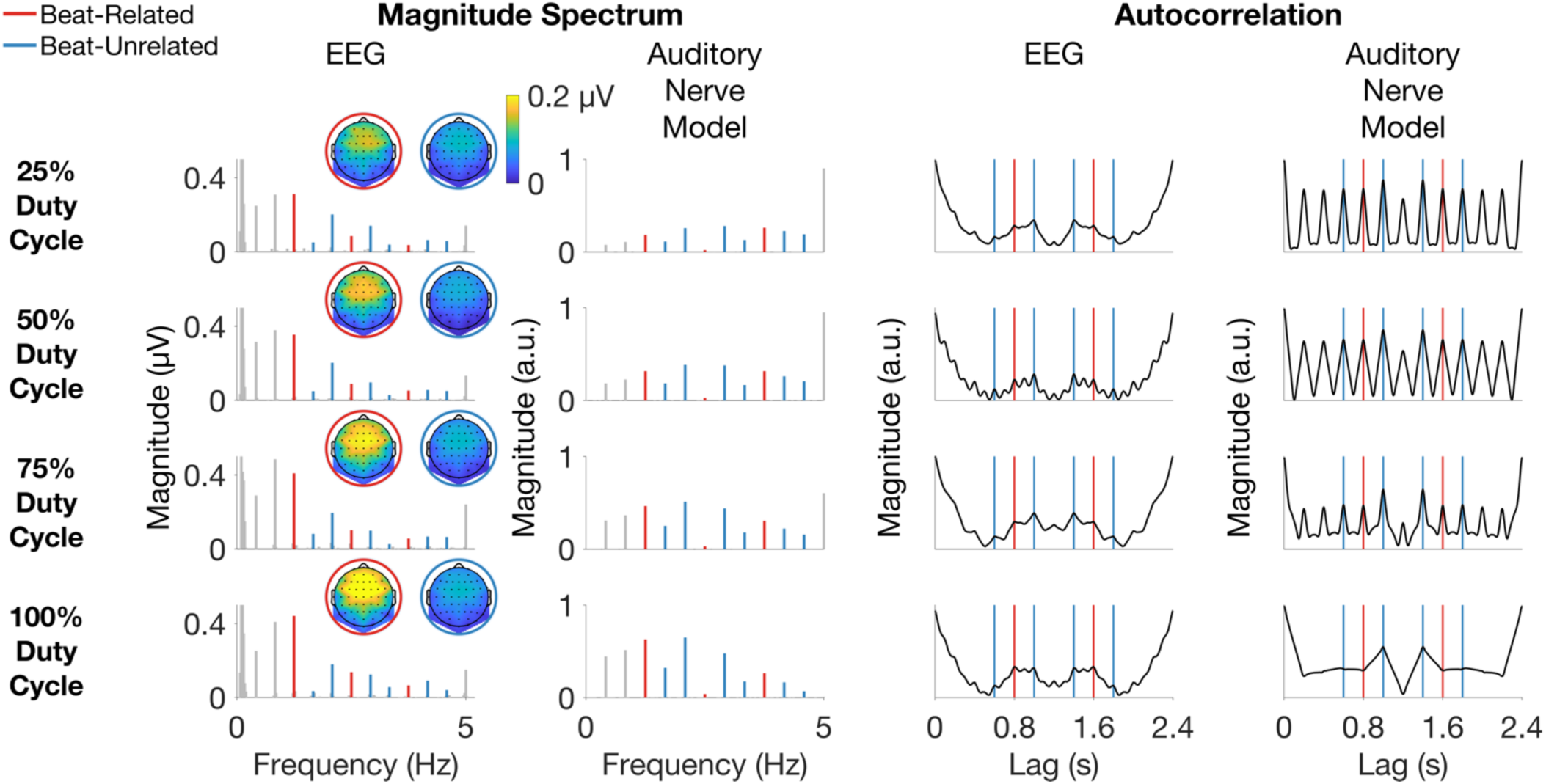
EEG and auditory nerve model output analysis based on magnitude spectrum and autocorrelation. Each row represents a duty cycle condition. The two columns on the left represent the magnitude spectrum-based analysis. The first column represents the group-level averaged magnitude spectra at a pool of fronto-central electrodes, across conditions. Beat-related frequencies are shown in red, and beat-unrelated frequencies are shown in blue. Scalp topographies of the neural activity measured at the average magnitudes of beat-related (in red circle) and unrelated (in blue circle) frequencies are represented as insets. The second column represents the normalized magnitude spectra obtained from the auditory nerve model output for each duty cycle sequence. The two columns on the right represent the autocorrelation-based analysis (for visualization purposes, only a subset of lags from 0 to 2.4 s corresponding to the pattern duration is shown). The first column represents the group-level averaged autocorrelation function measured from the same pool of fronto-central electrodes, across conditions. Beat-related lags are shown in red, and beat-unrelated lags are shown in blue. The second column represents the autocorrelation function of the auditory nerve model output for each duty cycle sequence.

**Figure 3.**
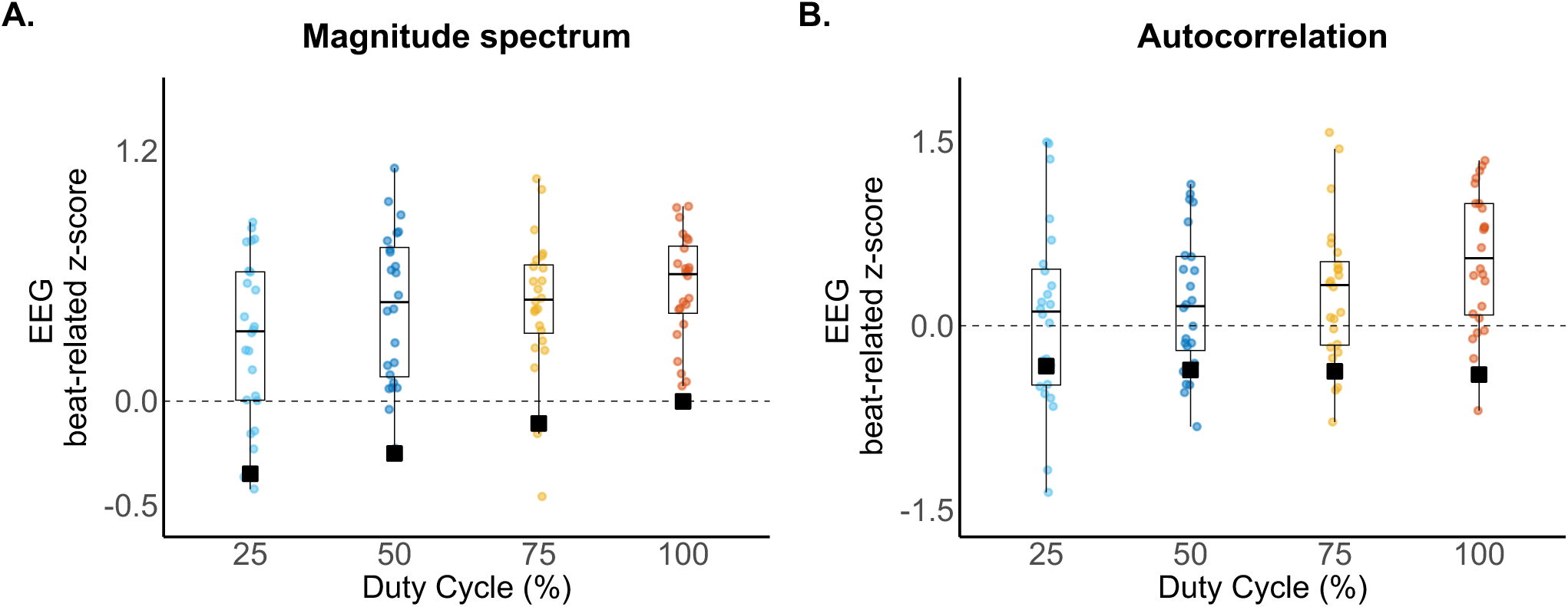
Increased representation of beat periodicities in the EEG with longer duty cycle. A) Magnitude spectrum-based analysis. For each duty cycle condition, beat-related z-scores measured from EEG responses of individual participants are shown as circles. The corresponding z-scores obtained from the auditory nerve model are depicted as black squares and were all inferior the EEG responses and to zero, reflecting non prominent beat periodicity at this peripheral stage of processing. Horizontal line, boxplot and whiskers indicate the median, 25th and 75th percentiles, and minimum and maximum values, respectively. B) Autocorrelation-based analysis. Same structure as panel A.

A significant main effect of duty cycle on the mean beat-related z-scores was observed (F₃,₆₉ = 3.44, p = 2.14 × 10⁻², η_p_^2^ = 0.13), indicating that beat-related EEG responses differed across duty cycle conditions. EMMs showed progressively increasing beat-related z-scores for longer duty cycles (0.30, 0.45, 0.47, 0.55, respectively).

Post-hoc pairwise comparisons revealed significantly greater beat prominence for the 100% compared with the 25% duty cycle condition (estimate = 0.25, SE = 0.08, t₆₉ = 3.13, p = 1.55 × 10⁻²), whereas no other pairwise comparisons reached significance after Bonferroni correction (see Table 1 for details). Post-hoc polynomial contrast demonstrated a significant positive linear trend across duty cycle levels (t₆₉ = 3.04, p = 3.30 × 10⁻³), confirming that beat-related neural responses increased systematically with increasing duty cycle.

**Table 1.**
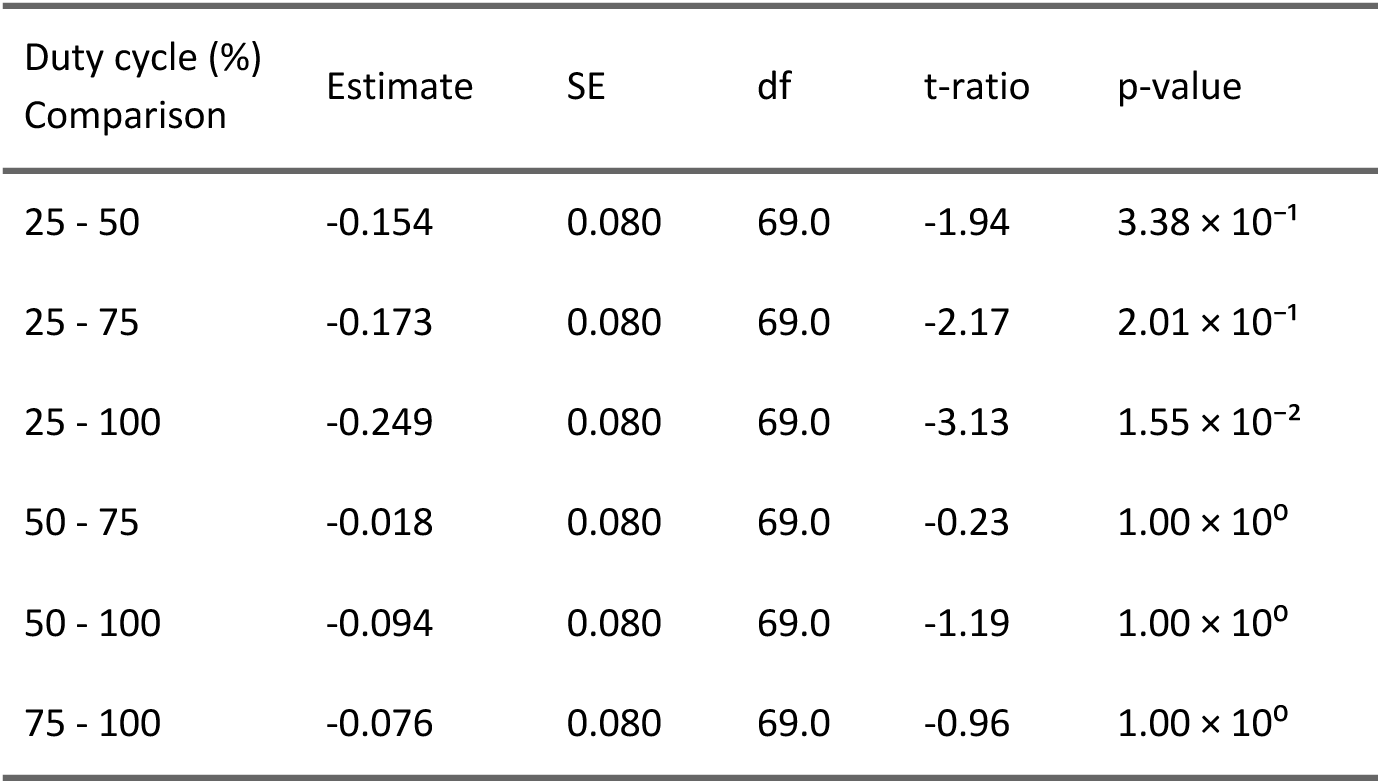
Magnitude spectrum-based analysis: post-hoc pairwise comparisons of EEG beat-related z-scores across duty cycle conditions.

| Duty cycle (%)<br>Comparison | Estimate | SE | df | t-ratio | p-value |
| --- | --- | --- | --- | --- | --- |
| 25 - 50 | -0.154 | 0.080 | 69.0 | -1.94 | $3.38 \times 10^{-1}$ |
| 25 - 75 | -0.173 | 0.080 | 69.0 | -2.17 | $2.01 \times 10^{-1}$ |
| 25 - 100 | -0.249 | 0.080 | 69.0 | -3.13 | $1.55 \times 10^{-2}$ |
| 50 - 75 | -0.018 | 0.080 | 69.0 | -0.23 | $1.00 \times 10^0$ |
| 50 - 100 | -0.094 | 0.080 | 69.0 | -1.19 | $1.00 \times 10^0$ |
| 75 - 100 | -0.076 | 0.080 | 69.0 | -0.96 | $1.00 \times 10^0$ |

In addition, auditory nerve model z-scores significantly predicted EEG z-scores across conditions (F₁,₇₁ = 8.86, p = 3.99 × 10⁻³, η_p_² = 0.11; estimate = 0.63, SE = 0.21). Yet, the EEG went beyond the auditory nerve model. That is, beat-related z-scores were significantly higher in the neural responses as compared to the auditory nerve model across all duty cycle conditions, indicating significant periodization of neural activity (all ps < 1 × 10⁻⁴; see Table 2 for details).

**Table 2.** Magnitude spectrum-based analysis: comparison of EEG beat-related z-scores (estimated marginal means, EMMs) against auditory nerve model z-scores for each duty cycle condition.

| Duty cycle (%) | EEG EMM | SE | df | AN model z-score | t-ratio | p-value |
| --- | --- | --- | --- | --- | --- | --- |
| 25 | 0.30 | 0.069 | 68.2 | -0.348 | 9.30 | $< 1 \times 10^{-4}$ |
| 50 | 0.45 | 0.069 | 68.2 | -0.250 | 10.12 | $< 1 \times 10^{-4}$ |
| 75 | 0.47 | 0.069 | 68.2 | -0.107 | 8.31 | $< 1 \times 10^{-4}$ |
| 100 | 0.55 | 0.069 | 68.2 | -0.001 | 7.89 | $< 1 \times 10^{-4}$ |

Finally, while the auditory nerve model outputs showed beat-related z-scores that were monotonically increasing with longer duty cycles, these z-scores remained negative across duty cycles (−3.48 × 10⁻¹, −2.50 × 10⁻¹, −1.07 × 10⁻¹ and −9.31 × 10⁻⁴, from shorter to longer duty cycle conditions, respectively), confirming that beat periodicities were not prominent in the acoustic inputs and their lower-level sensory representations in *any* of the four stimulus sequences.

### 3.2 Autocorrelation-based analysis

As with the magnitude spectrum-based approach, a significant main effect of duty cycle was also found with the autocorrelation-based approach on the EEG beat-related z-scores (F₃,₆₉ = 3.66, p = 1.65 × 10⁻², η_p_² = 0.14), with increasing EMMs of beat-related z-scores as the duty cycle increased (0.08, 0.18, 0.27, 0.54, respectively). Bonferroni-corrected post-hoc comparisons also revealed significantly greater beat prominence for the 100% compared to the 25% duty cycle condition (estimate = 0.47, SE = 0.15, t₆₉ = 3.16, p = 1.40 × 10⁻²), with no other pairwise contrasts reaching significance (see Table 3). A post-hoc polynomial contrast revealed a significant positive linear trend across conditions (t₆₉ = 3.19, p = 2.1 × 10⁻³), confirming stronger beat-related periodicity for longer duty cycles, as found with the magnitude spectrum-based approach.

**Table 3.**
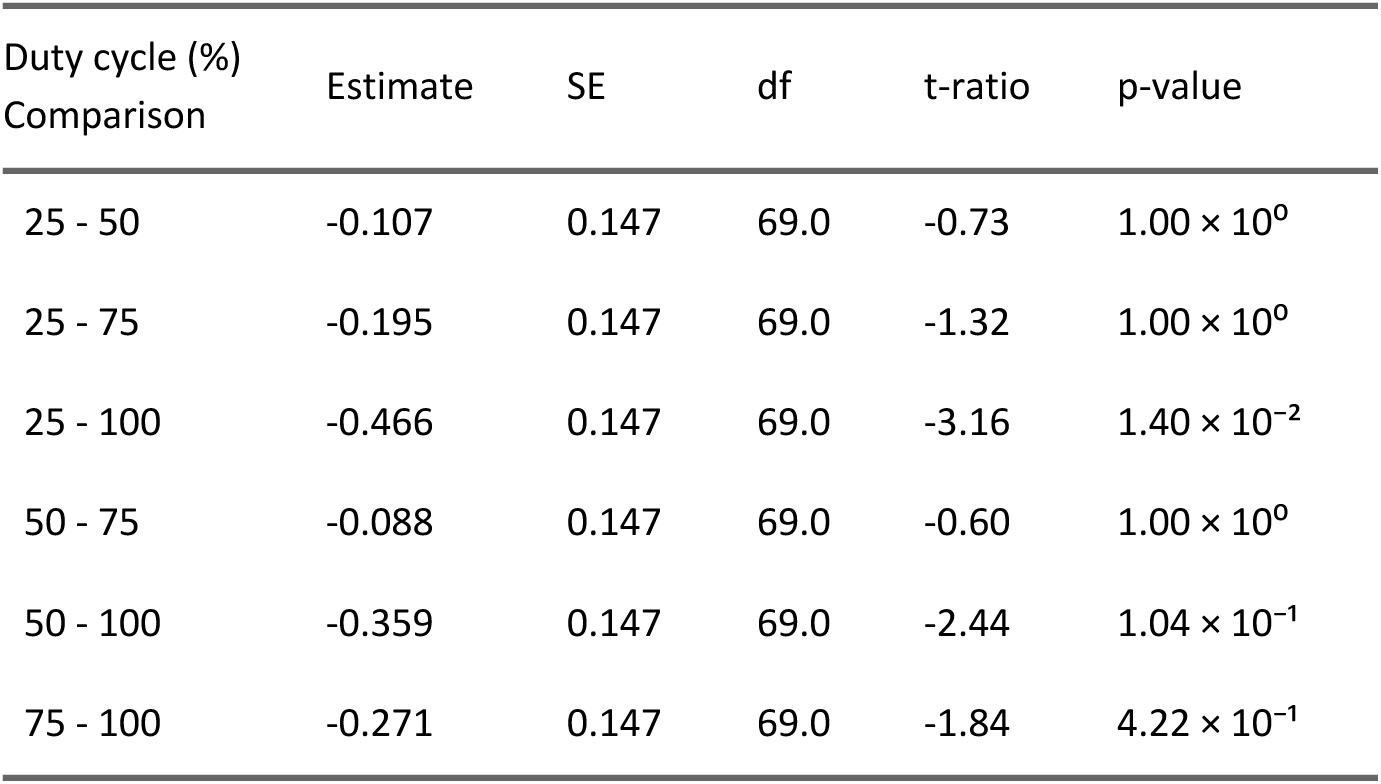
Autocorrelation-based analysis: post-hoc pairwise comparisons of EEG beat-related z-scores across duty cycle conditions.

In contrast with the increased neural representation of the beat with longer duty cycles, auditory nerve model z-scores were all negative and comparable across conditions (−3.30 × 10⁻¹, −3.61 × 10⁻¹, −3.72 × 10⁻¹, and −3.98 × 10⁻¹, respectively), confirming the lack of prominent beat periodicities in the auditory nerve model outputs. This relative stability across duty cycles reflects the fixed temporal structure of sound onsets across sequences, together with autocorrelation’s invariance to individual event shape when capturing beat periodicity. EEG beat-related z-scores were significantly greater than in the corresponding auditory nerve model for all duty cycle conditions, showing enhanced representation of beat periodicities in the EEG as compared to the lower-level auditory representations as found with the magnitude spectrum-based analysis (all p ≤ 2.1 × 10⁻³; see Table 4).

**Table 4.** Autocorrelation-based analysis: comparison of EEG beat-related z-scores (estimated marginal means, EMMs) against auditory nerve model z-scores for each duty cycle condition.

| Duty cycle (%) | EEG EMM | SE | df | AN model z-score | t-ratio | p-value |
| --- | --- | --- | --- | --- | --- | --- |
| 25 | 0.08 | 0.127 | 69.4 | -0.330 | 3.20 | $2.06 \times 10^{-3}$ |
| 50 | 0.18 | 0.127 | 69.4 | -0.361 | 4.29 | $< 1 \times 10^{-4}$ |
| 75 | 0.27 | 0.127 | 69.4 | -0.372 | 5.07 | $< 1 \times 10^{-4}$ |
| 100 | 0.54 | 0.127 | 69.4 | -0.398 | 7.41 | $< 1 \times 10^{-4}$ |

Together, these results thus indicate significant periodization of brain responses across all duty cycle conditions, with strongest beat-related responses for the longest duty cycles, irrespective of the waveform shape of the input or output. Importantly, the convergence of the results across both the magnitude spectrum-based and autocorrelation-based approaches allow us to rule out the contribution of nonspecific differences in response waveform shape across conditions. Instead, it supports the interpretation that increased beat-related z-scores with longer duty cycles, as well as the larger beat-related z-scores observed in EEG relative to the auditory nerve model, both reflect enhanced response periodicity.

### 3.3 Low-frequency range as a preferential bandwidth for neural responses to acoustic rhythms

In the auditory nerve model outputs, the proportion of magnitude concentrated in the low-frequency range (< 5 Hz) increased monotonically with duty cycle, yet without capturing a major part of the magnitude (30.71%, 44.09%, 45.37%, 48.92% of the total magnitude up to 45 Hz, see Table 5a for detailed values across frequency bands). In contrast, EEG group-level responses exhibited an overall magnitude predominantly concentrated within the low-frequency range (< 5 Hz) across all duty cycle conditions (77.22%, 67.02%, 75.45%, and 76.48% of the total magnitude up to 45 Hz; see Table 5b for detailed values across frequency bands). At the participant level, there was no evidence for a monotonic variation of the proportion of response magnitude below 5 Hz with duty cycle (F_1, 71_ = 0.70, p = 4.05 × 10⁻¹). Overall, these results identify the < 5 Hz range as the preferential frequency bandwidth for neural responses to rhythmic acoustic sequences, as it consistently captured a substantial proportion of the total EEG response magnitude irrespective of duty cycle.

**Table 5a.** Auditory nerve model outputs: proportion of total magnitude (0–45 Hz) in each 5-Hz frequency band, by duty cycle condition.

| Frequency band (Hz) | 25% | 50% | 75% | 100% |
| --- | --- | --- | --- | --- |
| 0–5 | 30.71% | 44.09% | 45.37% | 48.92% |
| 5–10 | 24.82% | 12.57% | 14.97% | 14.24% |
| 10–15 | 11.71% | 11.74% | 9.29% | 9.01% |
| 15–20 | 5.25% | 6.45% | 6.91% | 6.86% |
| 20–25 | 6.78% | 6.95% | 6.26% | 5.71% |
| 25–30 | 6.21% | 5.32% | 5.15% | 4.77% |
| 30–35 | 5.19% | 4.77% | 4.47% | 4.06% |
| 35–40 | 5.23% | 4.57% | 4.23% | 3.50% |
| 40–45 | 4.11% | 3.54% | 3.36% | 2.94% |

**Table 5b.** EEG group-level responses: proportion of total magnitude (0–45 Hz) in each 5-Hz frequency band, by duty cycle condition.

| Frequency band (Hz) | 25% | 50% | 75% | 100% |
| --- | --- | --- | --- | --- |
| 0–5 | 77.22% | 67.02% | 75.45% | 76.48% |
| 5–10 | 10.52% | 18.85% | 11.88% | 12.37% |
| 10–15 | 5.46% | 7.01% | 6.10% | 5.67% |
| 15–20 | 3.52% | 3.15% | 3.78% | 2.01% |
| 20–25 | 1.13% | 2.12% | 1.07% | 0.92% |
| 25–30 | 0.42% | 0.66% | 0.44% | 0.93% |
| 30–35 | 0.54% | 0.23% | 0.55% | 0.81% |
| 35–40 | 0.83% | 0.69% | 0.49% | 0.35% |
| 40–45 | 0.36% | 0.28% | 0.24% | 0.44% |

## 4 DISCUSSION

Understanding how sensory features shape neural representations of the beat is a key step toward elucidating the brain processes underlying rhythm perception. Here, we tested whether sound envelope duration (sonic duty cycle) modulates beat-related neural responses when the onset structure is held constant. We found that rhythms conveyed with longer sound durations elicited an enhanced neural representation of the perceived beat, although the rhythmic stimulus itself did not contain prominent periodicity corresponding to the perceived beat.

### A Holistic Processing Account of Beat Perception

Our study adds to a growing body of evidence suggesting that, beyond onset timing, other acoustic features such as pitch, rise time and sound duration also shape beat perception. Here, by manipulating sound duration while keeping onset timing and other acoustic parameters constant across conditions, we found that longer sound durations led to stronger neural activity at beat-related frequencies. These findings corroborate a holistic processing account of rhythm perception whereby neural representations of the beat rely on integration of the onset timing and other stimulus features such as the individual sound envelope duration.

The potential holistic nature of rhythm perception has important theoretical and methodological implications. For example, the type of stimulus features (e.g., sound duration) and their combinations (e.g., sound duration combined with rise time, pattern of onset timing and specific ranges of speed) may differentially facilitate association between a rhythmic stimulus and an internal representation of beat. In the present study, we observed that long duty cycle facilitated temporal integration of fast rhythmic onset timings into slower, behavior-relevant units experienced as the beat. Whether this facilitation emerges from hardwired non-linear processes of the human auditory system yielding temporal integration properties that are contingent on other stimulus parameters such as speed remains to be tested. Alternatively, this facilitation might be plastic, that is, expected to depend on the participant’s familiarity and experience with specific combinations of features that are statistically and socially prominent in their music-cultural environment ^21,23,86,87^.

### Revealing the Preferential Frequency Bandwidth Supporting Neural Representation of the Beat

Crucially, analyses of simulated auditory nerve model responses revealed a progressive redistribution of the magnitude toward the lower-frequency range (< 5 Hz) with increasing sonic duty cycle. This shift toward slower temporal fluctuations driven by stimulus envelope at the level of the auditory periphery may be expected to increase the overall gain of neural responses, in line with the low-pass characteristic of the auditory modulation transfer function. In contrast, we found EEG responses to be predominantly concentrated in the low-frequency range, and this was the case irrespective of the duty cycle. This observation aligns with previous work implicating low-frequency cortical activity as a critical bandwidth supporting auditory processes such as temporal prediction ^88,89^, as well as speech and music encoding ^90–96^.

Taken together, these findings suggest that the increased sound duration associated with longer duty cycles may optimize the alignment between the physical features conveying the rhythmic stimuli and the brain’s preferential frequency band for higher-level internal representation of rhythm. Such alignment may particularly benefit temporal scaffolding processes involved in mapping fast acoustic onsets onto an internal representation of slower, behavior-relevant units experienced as the beat ^36,97^.

### Onset Structure of Rhythmic Inputs and Lower-Level Auditory Processing Does Not Explain Neural Enhancement of Beat Periodicities

In the present study, the rhythmic stimuli used in all conditions were purposely designed to consistently elicit a beat perceived at a specific periodicity, while ruling out a lower-level confound by lacking prominent acoustic modulations at this periodicity ^25,50,51,55,56,72,78,79^. A biomimetic model of peripheral auditory processing confirmed the absence of prominent representation of the beat periodicity in the stimulus. The assumption that this stimulus would nonetheless elicit a specific beat percept was based on previous research using comparable (Western) participant groups ^25,36,50,51^. It was also confirmed in the present study through behavioral (tapping) responses to the stimuli (see Supplementary Materials).

Our finding of periodized neural activity during static listening therefore adds to the growing body of EEG research showing that the human brain is capable of (i) producing behavior-relevant periodized activity regardless of the prominence of these periodic features in the input, and (ii) this periodization emerges implicitly. These results therefore corroborate the notion that higher-level neural processes selectively boost the representation of the beat when it is not prominent in the lower-level early stages of processing ^15,18,25,49–51,56,72,77–79,98–100^. Moreover, our findings are also in line with behavioral and qualitative music research emphasizing that the beat can be experienced without being prominent in the acoustic signal ^15,101–105^. Taken together, our results thus reinforce the view of beat perception as a form of perceptual categorization, in which rhythmic inputs are mapped onto a "periodized" representation.

### Longer Sound Duration Leads to Larger Temporal Integration Window, Beyond Discrete Tracking of Individual Onsets

Auditory nerve modeling showed that shorter duty cycles yielded comparable population firing rates across successive sounds within a rhythmic group. In contrast, for the longest duty cycle, responses to the group’s first sound were enhanced relative to subsequent sounds, reflecting adaptation processes in the auditory periphery (see Fig. 1) ^106^.

Notably, such adaptation may, in certain cases, enhance the representation of the beat periodicity by emphasizing the grouping of sounds rather than individual sounds making up the rhythmic sequence ^107^. However, this would only happen in cases where the groups of tones are already arranged in a way that aligns with the beat (such rhythmic stimuli have been referred to in prior work as “unsyncopated”, “strongly periodic”, “strongly metrical”, etc.) _15,19,46,99 ._

In contrast, when the internal representation of the beat does not match the pattern of onset timings of the sensory input, the brain must go beyond these lower-level stages of processing. In such cases, one could speculate that longer duty cycles—stimulating the auditory system with an acoustic envelope particularly concentrated toward lower-frequency ranges (< 5 Hz)—foster integration of the fast onsets into slower units perceived as the beat, thus beyond discrete tracking of individual onsets. This interpretation is consistent with behavioral findings showing that longer sounds are associated with broader and delayed perceptual integration windows, or “beat bins”, relative to sound onset ^21^. Such expanded integration windows may allow the system to organize the internal representation of individual sound events in the rhythmic stimulus more flexibly, even beyond internal representations of groups of tones. This is particularly relevant in rhythms where the pattern of onset timings is not aligned with the beat periodicity (such as the rhythm used in the current study), and in line with the emerging understanding of how acoustic features are used in contemporary music practices to promote movement synchronization ^108–110^. In other words, longer sounds may activate mechanisms that would promote higher-level non-linear transformation of the input, further amplifying the brain’s capacity to go beyond a one-to-one mapping between stimulus features and the internal representation of the beat ^99^.

## 5 CONCLUSION

Our findings demonstrate that longer sound duration enhances neural representation of a periodic beat, even when those beat-related periodicities are not acoustically salient in the stimulus. This enhancement was associated with a shift of the input’s envelope toward the lower frequency range preferentially tracked by cortical responses to rhythmic inputs. Together, these results highlight a privileged bandwidth for beat processing and corroborate a holistic processing account of rhythm whereby onset timing and sound envelope duration are integrated to yield higher-level internal representations of rhythm.

## Acknowledgments

The work was supported by a FRIA-FNRS grant (F.R.) and Grants from the European Research Council (801872, 101228872, and 101148958).

## Authors’ contributions

F.R., C.L., T.L. and S.N. designed the study, F.R. performed the study, F.R., C.L., T.L., and S.N. analyzed data, created the figures and wrote the paper. All authors contributed to writing and editing the paper.

## Data availability statement

Anonymized raw data are available at the Open Science Framework (OSF). Link: (https://osf.io/mkg8d/overview?view_only=a102e8de0b834235aa3537d55819d8e3).The scripts used to conduct the analyses and generate the figures are available on a public GitHub repository (https://github.com/florarosen/Audio_DutyCycle).

## List of abbreviations

EEG: Electroencephalography
IOI: Inter-Onset Interval
RMS: Root Mean Square
SNR: Signal-to-Noise Ratio
FFT: Fast Fourier Transform
LMM: Linear Mixed Model
EMMs: Estimated Marginal Means
ICA: Independent Component Analysis

