## Supplementary Materials for "Beyond Onset Timing: Longer Sound Envelope Duration Enhances Neural Representation of the Musical Beat"

### 1. TAPPING EXPERIMENT AND ANALYSIS

Each EEG “static” block was followed by a "tapping" block of four trials, in which participants were instructed to tap along with the periodic beat they would perceive while listening to the same 40.8-s sequence as in the EEG blocks. These tapping blocks aimed to inform on the period of the beat perceived across conditions and participants. This allowed us to further determine whether each frequency or lag of interest would be categorized as beat-related (if matching the beat period and its harmonics or multiples) or not, in the main EEG analysis.

#### 1.1. Experimental Design

Prior to the experiment, participants completed a familiarization phase consisting of an unrecorded training session where we ensured that the tapping task was understood. During this familiarization phase, they were presented with three real-world musical excerpts and one rhythmic sequence composed of similar types of sounds but conveying a different rhythmic pattern as compared to the experimental stimuli. Participants were instructed to tap the space bar of a laptop keyboard as regularly as possible along with the beat they would perceive from the excerpts (instruction: "Tap along with the beat you naturally feel in the music, as if you were clapping your hands or nodding your head at a concert.”). After a few seconds, a superimposed clap track indicated an example plausible beat aligned to the excerpt, to illustrate the type of expected tapping responses (i.e., periodic, continuous over the excerpt, and synchronized with the sequence). It was emphasized that tapping at a different rate was not incorrect, but participants were instructed to synchronize with the claps once heard in order to try feeling the beat period being highlighted. Visual feedback of their accuracy was displayed on the screen until the end of each trial, to allow for online correction of the period and phase of taps to match the cued beat periodicity. After this familiarization phase, participants proceeded with the experiment.

Following ten “static” listening trials during which EEG was recorded, participants completed four “tapping” trials. In these, participants were instructed to tap the beat period they perceived with their preferred index finger using a custom-built response box (hereafter named "tapping box"). The tapping box contained a high resistance switch able to generate a trigger signal every time the fingertip contacted the response box (Institute of Neurosciences, UCLouvain, Belgium). The tap triggers were recorded through an extra Analog Input Box daisy chained to the AD-box of the Biosemi Active-Two system (Biosemi, Amsterdam, The Netherlands) used to record the EEG in the “static” sessions and digitized at a 1024 Hz sampling rate.

Participants were asked to wait no more than one or two repetitions of the pattern after the sequence had started before beginning to tap, and to tap as regularly as possible until the sequence would finish. It was emphasized that the goal was to feel as aligned with the sequence as possible, without any specific period being expected. During these tapping trials, participants were also allowed to move other parts of their body (e.g., head, torso, feet) and close their eyes if it helped them feel more into their perceived beat. If participants' tapping was reproducing each acoustic onset instead of an isochronous pulse, the trial and instructions were repeated.

### 1.2. Materials and Methods

Tapping data were preprocessed using the Letswave 6 toolbox (https://www.letswave.org/) and custom-made MATLAB scripts running under MATLAB R2020a. For each participant and condition, time series of tap onsets collected from the tapping box were segmented from +2.4 s to +40.8 s relative to the sequence onset, which yielded four 38.4 s epochs per participant and condition. The first 2.4 s were excluded in order to discard taps that may have occurred within the first presentation of the rhythmic pattern and account for variability in the start of the sensorimotor synchronization across participants and trials. Upon further visual inspection of individual time series, trials presenting technical issues were discarded (one trial discarded from one condition for three participants, and one trial discarded from two conditions for one participant).

#### *1.2.1. Inter-tap-intervals*

Inter-tap-intervals (ITIs) were calculated for all epochs as the time between two consecutive tap onsets. ITIs shorter than 210 ms were considered artifacts (as they would correspond to tapping rates close to the shortest IOIs in the rhythmic pattern, 200 ms) and were thus excluded from further analysis. Furthermore, to determine which beat periodicity participants had synchronized to, the median ITI was calculated separately for each trial, condition, and participant. Each median ITI was then compared to beat periodicities that were considered as theoretically plausible given the pattern grid structure (i.e., periodicities composed of an integer number of times the shortest IOI could fit within the rhythmic pattern duration, that is, 400, 600, 800, 1200, and 2400 ms, corresponding to the shortest IOI x2, x3, x4, x6 and x12, respectively). The beat period targeted by participants' tapping in each trial was determined as the theoretically plausible beat period showing the smallest absolute percent difference with the median ITI. The median ITI values averaged across trials for each participant were further compared across conditions.

The targeted beat at the group level obtained from this ITI analysis was considered to represent the perceived beat periodicity, thus further used to categorize the harmonics of the pattern repetition rate as beat-related or -unrelated in the main analysis of EEG magnitude spectrum (and, similarly, to categorize the lags as beat-related and -unrelated, in the EEG autocorrelation function).

#### *1.2.2. Measuring beat prominence in the tapping responses: Magnitude spectrum-based analysis*

For each participant and condition, the obtained four 38.4 s tapping epochs were transformed into the frequency domain using Fast Fourier Transform (FFT), yielding a normalized magnitude spectrum ranging from 0 to 512 Hz with a frequency resolution of 0.026 Hz (1/38.4 s). The resulting magnitude spectra were baseline corrected by subtracting at each bin the average magnitude of bins 2 to 11 on either side, excluding immediately neighboring bins. Spectra were then averaged across trials, resulting in a single noise-subtracted magnitude spectrum per condition and participant. The prominence of beat-related frequencies was then quantified in the same way as for the EEG analysis (main text). Specifically, magnitudes at the frequencies of interest were extracted, standardized as z-scores, and z-scores at beat-related frequencies were averaged. This beat-related z-score value reflects the prominence of the periodicity corresponding to the beat, relative to the prominence of the other periodicities composing the signal [1].

#### *1.2.3. Measuring beat prominence in the tapping responses: Autocorrelation-based analysis*

To further evaluate the relative prominence of beat periodicities in the tapping data, the autocorrelation-based analysis was applied to the time-domain epochs. The autocorrelation function was obtained using the same procedure as for the EEG. Values at lags of interest corresponding to the grid rate and its multiples (i.e., corresponding to the shortest inter-onset interval in the sequences and its multiples) were then extracted, standardized as z-scores, and values at beat-related lags (800 ms and its multiples) were averaged to quantify the relative prominence of beat periodicity vs. other periodicities in the tapping responses.

### **1.3. Statistical analysis**

#### *1.3.1. Median ITI*

To compare the period of the tapped beat across duty cycle conditions, the median ITI was first averaged across all trials, separately for each condition and participant. Normality was assessed using the Shapiro-Wilk test, and homogeneity of variance was examined with Levene's test from the 'car' package in R (version 3.0-5). Differences across conditions were assessed using a non-parametric Friedman rank-sum test, with Kendall's W computed to measure agreement.

#### *1.3.2. Measuring beat prominence in the tapping responses across duty cycle conditions.*

For both the magnitude spectrum-based and the autocorrelation-based analyses pipelines, the prominence of the beat periodicity in the behavioral responses relative to its representation in the auditory nerve model was assessed using Wilcoxon signed-rank tests. For each condition, tapping z-scores were compared with the corresponding auditory nerve model output beat-related z-scores. The effect of duty cycle condition on beat prominence was further evaluated using non-parametric Friedman tests as normality assumptions were violated.

### 1.4. Results

#### *1.4.1. Converging median ITI values across duty cycle conditions*

The obtained median ITI values were similar across conditions, converging toward 800 ms, with narrow interquartile ranges (IQR), suggesting a tight clustering of ITI values around the median (see Figure S1A). A non-parametric Friedman rank sum test yielded no significant differences across conditions ($\chi^2(3) = 1.87$, $p = 6.00 \times 10^{-1}$, Kendall's $W = 0.03$).

#### *1.4.2. Significantly prominent beat periodicity in behavior across all duty cycle conditions*

When comparing tapping z-scores with auditory nerve model z-scores, all comparisons revealed significant differences ($ps < 1.00 \times 10^{-3}$) for both magnitude spectrum-based and autocorrelation-based analyses, suggesting that tapping responses were significantly more periodic than expected from the lower-level auditory representations across all conditions (see Figure S1B and C). No significant differences across duty cycle conditions were found for both magnitude spectrum-based z-scores ($\chi^2(3) = 4.05$, $p = 0.26$, Kendall's $W = 0.056$), and autocorrelation-based z-scores ($\chi^2(3) = 2.45$, $p = 0.48$, Kendall's $W = 0.034$).

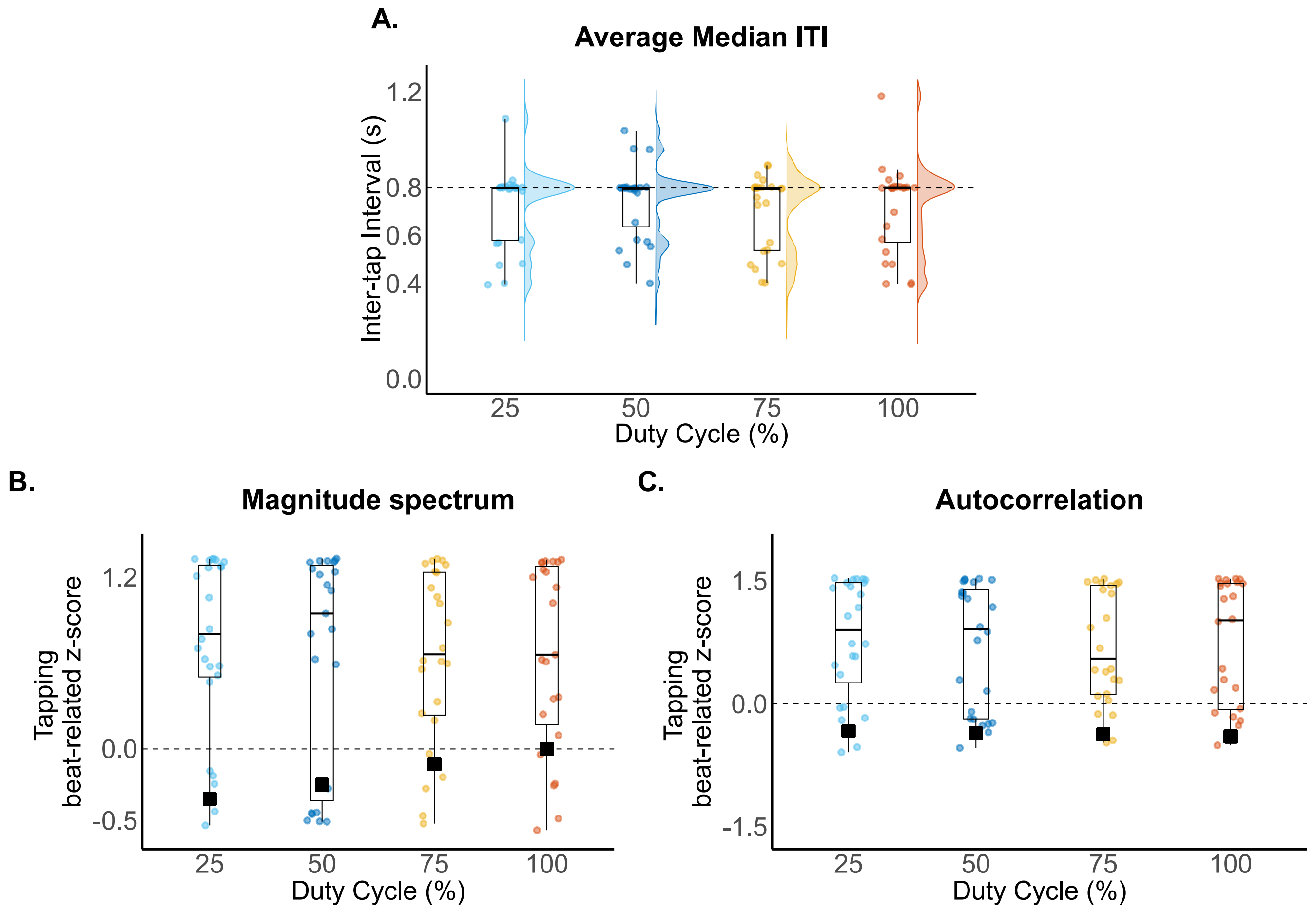


**Figure S1. Tapping analysis showing convergent beat period corresponding to four times the shortest IOI, and no significant differences in tapped beat across conditions.**

A) Inter-tap Interval (ITI). Each circle represents the median ITI for each participant and duty cycle condition. The dotted line represents the beat period of 0.8 s. B) Magnitude spectrum-based analysis. For each duty cycle condition, colored individual circles represent the average tapping beat-related z-score for each participant (horizontal line, boxplot and whiskers indicating the median, 25th and 75th percentiles, and min. and max. values, respectively). C) Autocorrelation-based analysis. Same structure as panel B.

### 2. MAGNITUDE SPECTRUM–BASED EEG ANALYSIS EXTENDED UP TO 15 HZ

#### 2.1. Materials and Methods

In the main text, analyses were restricted to the harmonics of the rhythmic pattern repetition rate located within a low-frequency range (<5 Hz, i.e., up to the shortest IOI). This frequency range was determined based on the distribution of amplitude observed in the EEG spectra, showing major part of the neural response concentrated below 5 Hz, in line with prior EEG studies using similar stimuli and tempo (see for example [2,3]).

To assess whether the observed effects of sonic duty cycle on beat-related neural responses depended on the frequency bandwidth selected for analysis, we performed a control EEG analysis on a larger range of harmonics of the rhythmic pattern rate, up to 15 Hz. Indeed, while the <5 Hz bandwidth encompassed a major portion of the response amplitude (as reported in the main text), the broader <15 Hz range captured nearly the totality of the neural response amplitude across all conditions (from shortest to longest duty cycle condition: 93.2%, 92.8%, 93.5%, 94.6%, based on group-averaged EEG magnitude spectra; see main Methods section for the details on this analysis). Analysis of the EEG responses and the auditory nerve model were identical to those described in the main Methods, with the harmonics of the rhythmic pattern repetition rate ranging up to 15 Hz.

#### 2.2. Results

*2.2.1. Beat prominence increases with longer duty cycles*

The results obtained with the extended frequency range were all similar to the results obtained with the smaller frequency range. Specifically, the analysis confirmed a significant main effect of duty cycle on the beat-related z-scores ($F_{3,69} = 3.09$, $p = 3.28 \times 10^{-2}$, $\eta_p^2 = 0.12$), indicating that beat-related EEG responses differed across duty cycle conditions. Estimated marginal means (EMMs) showed a monotonic increase in beat-related z-scores with increasing duty cycle, indicating stronger beat prominence in neural responses to longer duty cycles (0.17, 0.25, 0.26, 0.31, from shortest to longest, respectively). Moreover, Bonferroni-corrected post-hoc comparisons indicated significantly greater beat-related z-scores for the 100% compared with the 25% duty cycle condition ($p = 2.4 \times 10^{-2}$), whereas no other pairwise contrasts reached significance after correction. Finally**,** post-hoc polynomial contrast showed a significant positive linear trend across duty cycle conditions ($t_{69} = 2.94$, $p = 4.5 \times 10^{-3}$), indicating that beat-related z-scores increased linearly with duty cycle.

#### 2.2.2. *Significant periodization of EEG responses across duty cycle conditions*

The analysis of the auditory nerve model outputs using the extended frequency range corroborated the main Results. Specifically, the control analysis confirmed that beat periodicities were not prominent in any condition in the lower-level sensory representations, although the corresponding beat-related z-scores increased monotonically with longer duty cycles **(**$-1.95 \times 10^{-1}$**,** $-6.76 \times 10^{-2}$**,** $-2.17 \times 10^{-2}$, and $2.80 \times 10^{-2}$**,** from shortest to longest duty cycle, respectively). Moreover, these auditory nerve model z-scores significantly predicted EEG z-scores across conditions ($F_{1,71} = 9.42$, $p = 3.04 \times 10^{-3}$, $\eta p^2 = 0.12$; estimate = 0.60, SE = 0.20). The control analysis also confirmed significantly greater EEG z-scores as compared to the corresponding auditory nerve model z-scores for each condition (all ps $< 1 \times 10^{-4}$), indicating that neural responses exhibited robust beat-related periodization compared to the lower-level representations.

Overall, extending the frequency range of the magnitude spectrum–based analysis up to 15 Hz yielded results consistent with the analysis restricted to the <5 Hz range, without strengthening effect size, while confirming a robust effect of the duty cycle. Moreover, overall, beat-related z-scores were numerically reduced for both EEG and auditory nerve model when considering the larger frequency range, supporting the idea that including harmonics with magnitude near the noise floor, i.e., from frequency ranges contributing minimally to the total magnitude of the response, merely deteriorated the measures, with no additional information beyond the low-frequency range. Together, these results further support the focus on the <5 Hz frequency range as an empirically relevant bandwidth for quantifying beat-related neural responses.

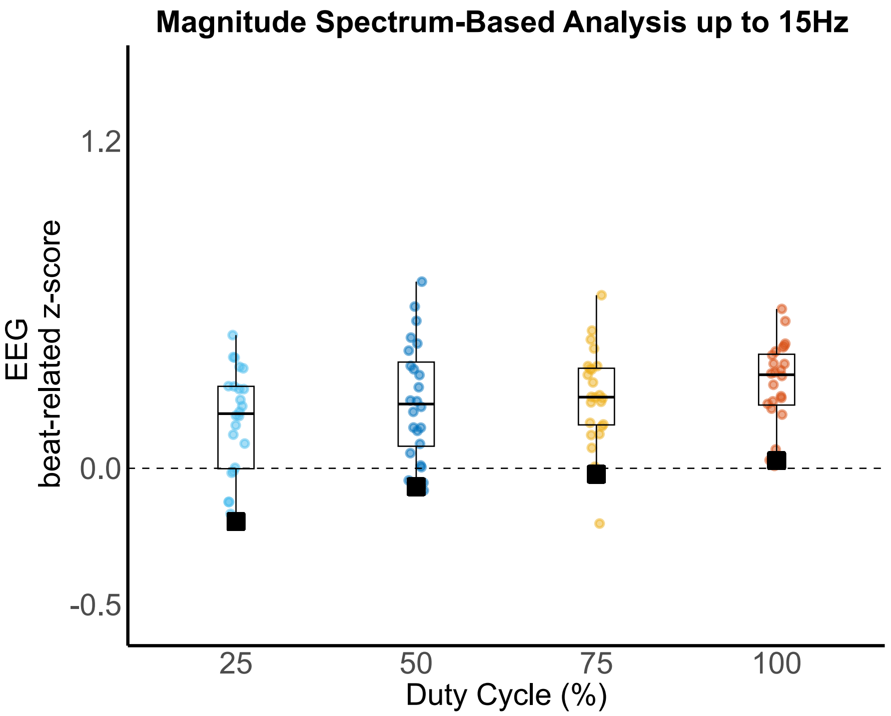


**Figure S2.** Magnitude spectrum-based analysis extended up to 15 Hz. Circles represent the EEG beat-related z-scores of individual participants for each duty cycle condition. The corresponding z-scores obtained from the auditory nerve model are depicted as black squares and were all equal or inferior to zero. Analysis revealed a significant effect of duty cycle condition on EEG z-scores, with significant positive linear trend across conditions indicating progressively stronger beat-related responses for longer duty cycles.
